# Predictive coding with spiking neurons and feedforward gist signalling

**DOI:** 10.1101/2023.04.03.535317

**Authors:** Kwangjun Lee, Shirin Dora, Jorge F. Mejias, Sander M. Bohte, Cyriel M.A. Pennartz

## Abstract

Predictive coding (PC) is an influential theory in neuroscience, which suggests the existence of a cortical architecture that is constantly generating and updating predictive representations of sensory inputs. Owing to its hierarchical and generative nature, PC has inspired many computational models of perception in the literature. However, the biological plausibility of existing models has not been sufficiently explored due to their use of artificial neural network features such as a non-linear, continuous, and clock-driven function approximator as basic unit of computation. Therefore, we have developed a spiking neural network for predictive coding (SNN-PC), in which neurons communicate using event-driven and asynchronous spikes. While adopting the hierarchical structure and Hebbian learning algorithms from previous PC neural network models, SNN-PC introduces two novel features: 1) a fast feedforward sweep from the input to higher areas, which generates a spatially reduced and abstract representation of input (i.e., a neural code for the gist of a scene) and provides a neurobiological alternative to an arbitrary choice of priors; and 2) a separation of positive and negative error-computing neurons, which counters the biological implausibility of a bi-directional error neuron with a very high basal firing rate. After training with the MNIST handwritten digit dataset, SNN-PC developed hierarchical internal representations and was able to reconstruct samples it had not seen during training. SNN-PC suggests biologically plausible mechanisms by which the brain may perform perceptual inference and learning in an unsupervised manner. In addition, it may be used in neuromorphic applications that can utilize its energy-efficient, event-driven, local learning, and parallel information processing nature.

**Author summary:** How does the brain seamlessly perceive the world, in the midst of chaotic sensory barrage? Rather than passively relaying information that sensory organs pick up from the external world along the cortical hierarchy for a series of feature extractions, it actively gathers statistical regularities from sensory inputs to track causal relationships between physical properties of external objects and the body. In other words, the brain’s perceptual apparatus is constantly trying to make sense of the incoming streams of sensory input and represent the subject’s current situation by building and maintaining internal models of the world and body. While this constructivist theme in understanding perception has been pervasive across multiple disciplines from philosophy to psychology to computer science, a comprehensive theory of brain function called predictive coding aims at unifying neural implementations of perception. In this study, we present a biologically plausible neural network for predictive coding that uses spiking neurons, Hebbian learning, and a feedforward visual pathway to perform perceptual inference and learning on images. Not only does the model show that predictive coding is well behaved under the biological constraint of spiking neurons, but it also provides deep learning and neuromorphic communities with novel paradigms of learning and computational architectures inspired by the nature’s most intelligent system, the brain.

## Introduction

In the midst of chaotic barrages of sensory information, the brain achieves seamless perception of the world. Despite the ease with which the brain achieves such a formidable feat, the problem of perception is computationally difficult, given that the brain has no direct access to the world. This renders perception into an inverse problem [1,2], in which the brain has to infer the distal stimulus in the physical world (i.e., cause) from the proximal sensations in the brain (i.e., effect) [3]. Moreover, given inherently noisy and ambiguous sensory information, the problem also becomes ill-posed. For example, exponentially many combinations of object arrangements and viewing conditions in the three-dimensional world can form the same two-dimensional retinal image.

How does the brain overcome such ambiguity, find a unique and stable solution to the inverse problem, and facilitate seamless perception of the world? A confluence of constructivist theories of perception [4–9] suggest that the brain imposes a priori constraints on possible solutions to the inverse problem based on an internal model of the world shaped by prior knowledge, experience, and context. In light of recent neurophysiological evidence that supports interaction of feedforward sensory inputs and feedback of a priori knowledge [10–13], predictive coding (PC) has been proposed as a possible neural implementation of perception [14–18]. According to PC in its canonical version [16], the brain employs hierarchical cortical circuits in which feedback connections carry predictions to lower areas, whereas feedforward connections carry the mismatch between actual and predicted neural activity (i.e., prediction error). The prediction error is used iteratively to correct the internal generative model, allowing to make more accurate inferences, but also to learn from errors. While its computational goal to explain away incoming sensory input resembles the ideas of redundancy reduction from information theory [19] and the efficient coding hypothesis [20], the probabilistic formalization of PC’s cortical processing algorithm that approximates Bayesian inference [21] builds on the Bayesian brain hypothesis [22] and Helmholtz machine [23]. In summary, PC offers a Bayes-inspired solution to the inverse and ill-posed problem of perception, and learning thereof, under the imperative of prediction error minimization.

Thanks to its potential to explain a multitude of cognitive and neural phenomena [14,16,24–29], PC has inspired many theoretical and computational models of perception. On the one hand, there are biologically motivated PC models in the literature to explain neural mechanisms of perception; on the other hand, machine learning inspired models seek missing ingredients that can place the perceptual capacity of artificial intelligence on par with the nature’s most intelligent machine. However, both approaches lack biological plausibility in their own respect. The biologically motivated PC models demonstrate how PC accounts for neuronal responses such as classical and extra-classical receptive field properties, but whether their efforts can be generalized across the cortical processing hierarchy remains an open question as their models were confined to specific components of the nervous system, such as the retina [14,24], the lateral geniculate nucleus [27], or V1 [26], or had a limited depth of processing hierarchy [16,26,28]. The machine learning inspired models show remarkable object recognition capabilities but lack the biological plausibility due to their reliance on supervised learning, convolutional filters, and backpropagation of errors [30–34]. Meanwhile, there has been an effort to bridge the gap between the two approaches: a deep gated Hebbian PC [35] successfully learns internal representations of natural images across multiple layers of the visual processing hierarchy, while exhibiting neuronal response properties such as orientation selectivity, object selectivity, and sparseness. Yet, previous models relied on an artificial neural network, the basic computational unit of which mimics a real neuron with limited biological realism [36] and communicates using synchronous and continuous signals instead of spikes.

To advance the biological realism of computational models of PC and move towards a more biologically plausible model of perception, we developed a spiking neural network for predictive coding (SNN-PC) by introducing two novel features: 1) a spiking neuron model [36,37] that describes the behavior of neurons with more biological details than firing-rate based artificial neurons, such as using binary, asynchronous spikes for synaptic communication and replacing a simple non-linear activation function with elaborate synaptic and membrane potential dynamics; and 2) a feedforward gist (FFG) pathway that mimics the gist of a scene or image [38], inspired by how the biological system of vision may rapidly recognize objects using a fast feedforward visual pathway [39–41]. While incorporating detailed biological mechanisms may lead to enhanced functionality, the resulting model turns out to lack a straightforward computational algorithm that accounts for negative error, which may arise from comparison between actual and predicted activities in PC circuits. In the following sections, we describe non-trivial problems in implementing a spiking version of deep predictive coding networks and offer our solutions. We show that, by putting together all the pieces, SNN-PC can learn hierarchical representations of MNIST hand -written digit images and infer unseen samples from spike signals of sensory inputs in a biologically plausible, unsupervised manner.

## Methods

The following section is organized into four subsections, which address challenges of implementing a PC neural network with spiking neurons and propose biologically plausible mechanisms to facilitate perceptual inference and learning: 1) introduction of a spiking neuron model; 2) description of synaptic communication between spiking neurons for reliable signal transmission and Hebbian learning; 3) separation of error-computing neurons into two groups to encode signed signals with dynamic binary spikes; and 4) introduction of the FFG pathway to establish informed initial conditions for prediction-generating neurons as opposed to random initialization.

### Single neuron model

The behavior of single neurons in SNN-PC was defined by the adaptive exponential integrate-and-fire model [42]:

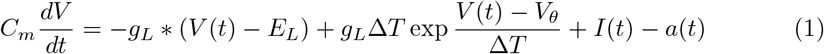

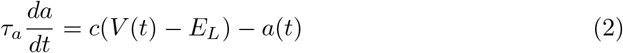

where *C_m_* is the membrane capacitance, *V*(*t*) is the membrane potential, *g_L_* is the leak conductance, *E_L_* is the leak reversal potential, Δ*T* is the slope factor, *V_θ_* is the action potential threshold, *a*(*t*) is the adaptation variable, *I*(*t*) is the incoming synaptic current, *τ_a_* is the adaptation time constant, and c is the adaptation coupling parameter. The parameter values are taken from [42] and listed in Table 1.

**Table 1.** Parameters for the adaptive exponential integrate-and-fire model

| Parameter | Value | Unit |
| --- | --- | --- |
| $C_m$ | 281 | $pF$ |
| $g_L$ | 30 | $nS$ |
| $E_L$ | -70.6 | $mV$ |
| $V_\theta$ | -50.4 | $mV$ |
| $\Delta T$ | 2 | $mV$ |
| $t_{ref}$ | 2 | $ms$ |
| $c$ | 4 | $nS$ |
| $b$ | 0.0805 | $nA$ |
| $\tau_a$ | 144 | $ms$ |
| $\tau_{rise}$ | 5 | $ms$ |
| $\tau_{decay}$ | 50 | $ms$ |

The membrane potential dynamics (Eq. 1) is described by a linear leak, a voltage-dependent exponential activation, which instantiates the fast activation of sodium channels [43], the incoming synaptic current, *I*(*t*), and an abstract adaptation variable, a, which couples the membrane potential dynamics with voltage-dependent subthreshold and spike-triggered adaptation (Eq. 2) [44].

At each time point of simulation, *t*, a neuron sums up all incoming current, *I*(*t*), at its postsynaptic terminal to update its membrane potential, *V*(*t*). Upon reaching the threshold, *V_θ_*, a spike is generated (i.e., *s*(*t*) = 1):

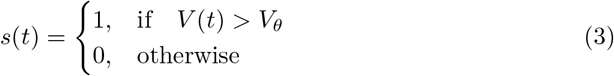

A spike is followed by an instantaneous reset of the membrane potential, *V*, to the resting potential, *V_r_*, and an increase of adaptation variable (*a*) by an amount, *b*, to model membrane potential repolarization and spike-triggered current adaptation, respectively:

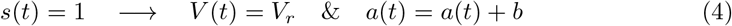

### Synaptic communication between spiking neurons

The behavior of the single neuron model described in the previous section (Eq. 1) is a function that takes incoming current, *I*(*t*), as input and a spike train, *s*(*t*), as output. For synaptic communication between spiking neurons, there must be a way to convert the output of a source neuron to input of a target neuron. The current entering a postsynaptic cell *j* (postsynaptic current; PSC) through *N* synapses from presynaptic cells *i* can be formulated as a continuous variable, *I_j_* (*t*), by applying an exponential low-pass filter to each incoming binary spike train to compute a spike trace, *X_i_*(*t*), and weighting each spike trace with the corresponding synaptic strength, *W_i,j_*, and summing the weighted spike traces over *N* synapses:

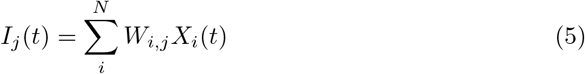

The spike trace, *X_i_*(*t*), is similar to the trace variable commonly used in spike timing dependent plasticity (STDP). With a proper choice of time constants (*τ_rise_* = 5 ms and *τ_decay_* = 50ms), it can be interpreted as NMDA receptor mediated synaptic transmission [45, 46]:

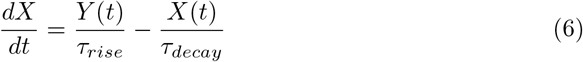

In particular, an NMDA-mediated component of PSC (‘NMDA current’) can be linked to the intracellular calcium concentration at the postsynaptic site, a high value of which leads to long term potentiation (LTP; [47,48]). Using the NMDA currents of pre- and postsynaptic neurons, synaptic weights are adjusted in the form of Hebbian learning. The learning of weights is described in detail in a subsequent section.

The first term in Eq. 6 governs the upward impact of the NMDA current corresponding to the influx of calcium into the postsynaptic neuron, whereas the second term describes the decay. The instantaneous reset of the variable, *Y*(*t*), that governs the increasing calcium influx resulting from the NMDA current following a presynaptic spike initializes glutamate release into the synaptic cleft, where it binds to NMDA receptors to open up calcium channels in the postsynaptic membrane:

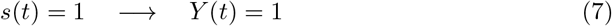

The glutamate concentration in the synaptic cleft decays exponentially back to the resting state (i.e., *Y*(*t*) → 0):

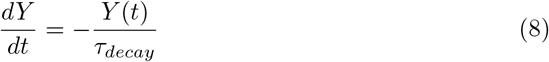

In summary, the synaptic communication consists of three major steps, which can be seen as a serial adaptation of the spike emission and reception filters in the spike response model [44]. The weighted spike trace (*W_i,j_ X_i_*(*t*)) approximates NMDA receptor mediated postsynaptic calcium transients. For example, consider the following case where presynaptic neurons (indexed by *i)* project to a postsynaptic neuron *j* (Fig 1):

**Fig 1.**
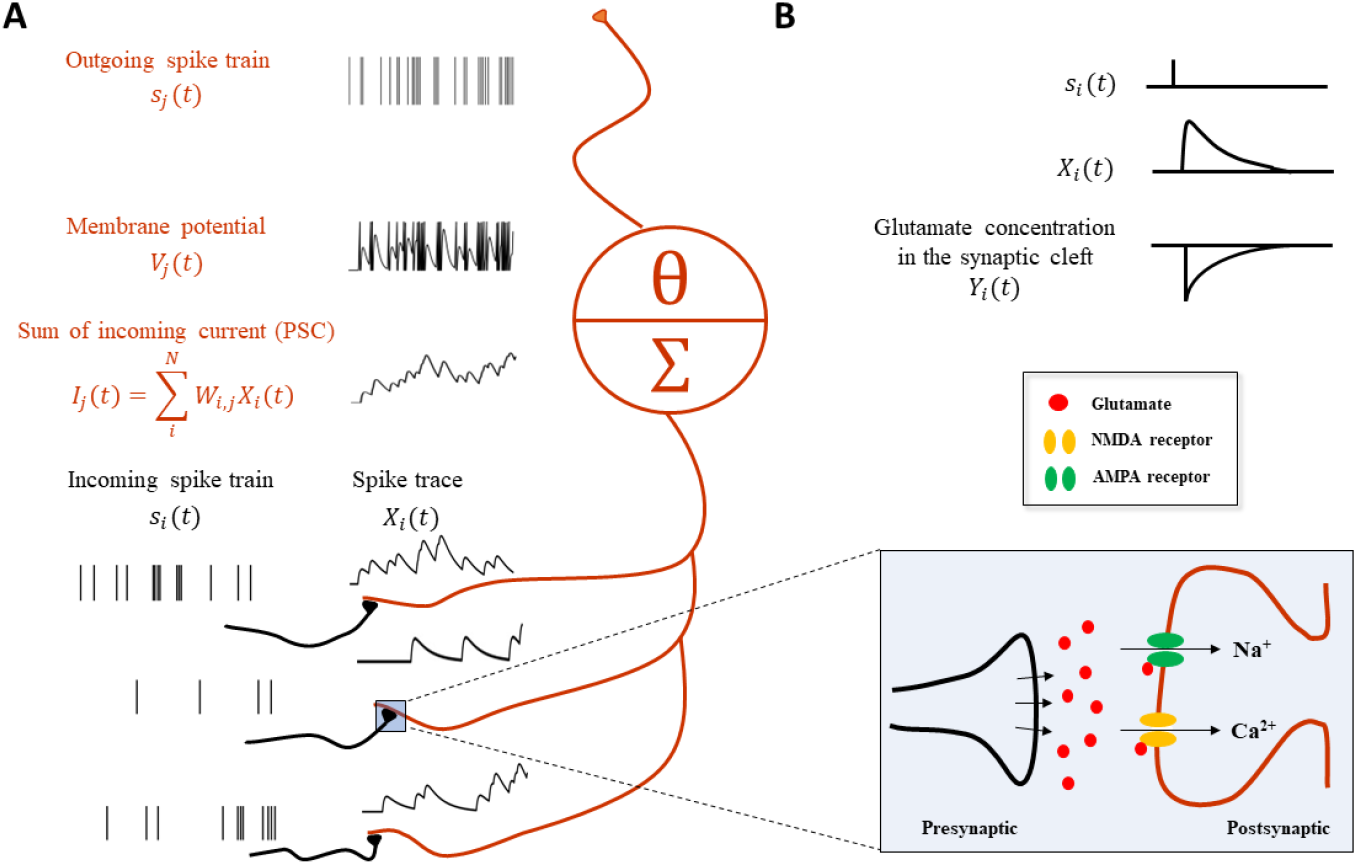
Synaptic transmission in a spiking neural network for predictive coding. (A) Schematic showing how spiking neurons in SNN-PC communicate with each other using spikes. Spikes from neurons *i* (black), *s_i_*(*t*), activating synapses that impinge on dendrites of neuron *j* (orange), are converted to spike traces, *X_i_*(*t*). The sum of all spike traces, weighted by synaptic strength, *W_i,j_*, make up the NMDA-mediated component of postsynaptic current, *I_j_*(*t*). In this particular scheme, we assume that all weights are 1 for simplicity. The postsynaptic membrane potential, *V_j_*(*t*), changes according to the incoming current and the cell emits spikes, *s_j_*(*t*), whenever it reaches threshold, *V_θ_*. (B) A presynaptic spike, *s_i_*(*t*) = 1, triggers glutamate release to the synaptic cleft (*Y*(*t*) = 1). When glutamate binds to the postsynaptic AMPA and NMDA receptors, the inward current of cations (*Na*^+^ and *Ca*^2+^) depolarizes the postsynaptic cell (*K*^+^ efflux not shown here for brevity). Concentrations of glutamate in the synaptic cleft and calcium in the postsynaptic terminal decrease exponentially (*Y_i_*(*t*) → 0 and *X_i_*(*t*) → 0, respectively) with time constants, *τ_rise_* and *τ_decay_*, respectively.

First, a spike train from neuron *i*, *s_i_*(*t*), is converted to an NMDA-mediated component of PSC received by neuron *j*, *I_j_*(*t*):

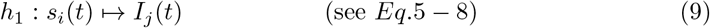

Second, the postsynaptic NMDA current arriving at the postsynaptic receptor site of neuron *j*, *I_j_*(*t*), influences the membrane potential of neuron *j*, *V_j_*(*t*):

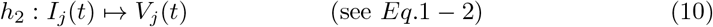

Third, the membrane potential, *V_j_*(*t*), generates a spike, *s_j_*(*t*), when it crosses the threshold, *V_θ_*:

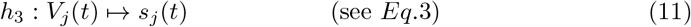

### Implementation of predictive coding

SNN-PC employed the same hierarchical structure as its two predecessors (Fig 2A) [16, 35]. Each area (denoted by superscript *l*) consists of two types of computational units (Fig 2): 1) a representation unit, 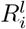, which infers the causes (i.e., generates latent representations) of incoming sensory inputs in the area below (*R*^*l*–1^) and makes predictions about the neural activity in the area below; and 2) an error unit, 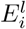, which compares the prediction from the area above with inputs from the area below and propagates the difference (i.e., prediction error) to the representation units in the area above (*R*^*l*+1^) to update the inferred causes and refine the prediction.

**Fig 2.**
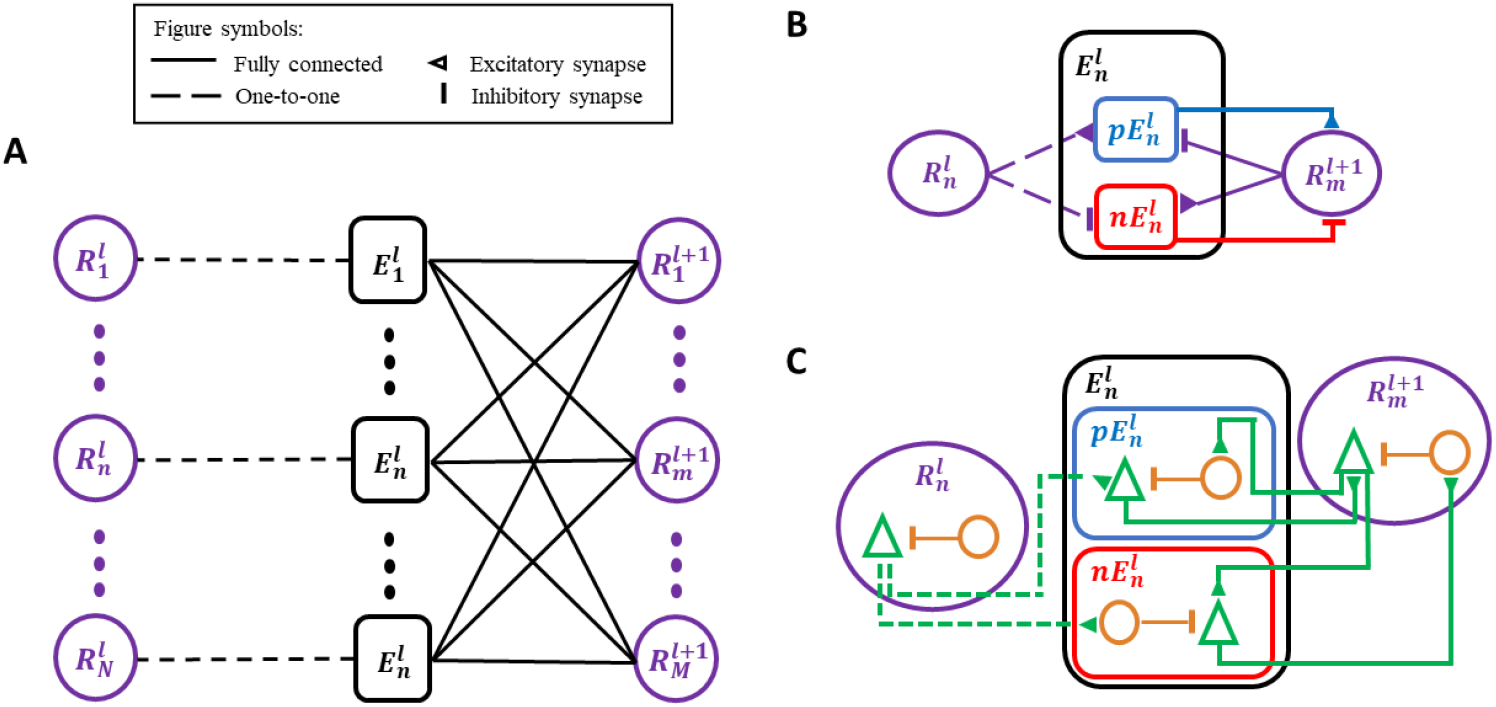
Adaptations of the classic predictive coding (PC) architecture to different levels of biological detail. (A) The classic PC architecture with representation (prediction) and error neurons as described by Rao and Ballard [16]. (B) To encode signed error signals with binary spikes, an error unit in *A*, 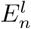, is separated into two units, 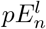 and 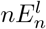, which compute a positive and negative error, respectively, via the opposite arrangement of excitatory-inhibitory synapses. The superscript (*l*) denotes the cortical processing area, whereas the subscript (*n* or *m*) denotes an index for unit in an area. (C) A biological interpretation of a computational unit in SNN-PC, which consists of a pyramidal cell (green triangle) and an interneuron (orange circle). An excitatory synapse exists between pyramidal cells of two units, whereas a polysynaptic inhibitory connection is formed by a pyramidal cell in a computational unit to an interneuron in another unit, which inhibits the pyramidal cell within the same unit.

#### Error unit

An error unit computes prediction errors by taking the difference between the sensory input in the lowest area, or its latent representation in the case of higher areas, and the corresponding prediction from the area above. Depending on the relative strengths of the two signals, this difference can be positive (i.e., input > prediction) or negative (i.e., prediction < input). The signed nature of the prediction error poses no obstacle to PC models with artificial neurons, which can encode positive and negative signals. However, given the non-negative nature of spike signals, SNN-PC has to adopt a different solution that can encode both types of prediction error. As observed in the dopaminergic system, a neuron may encode both types of prediction error by expressing the magnitude of error in relation to its basal firing rate: positive errors are encoded with activity above its basal firing rate and negative errors with activity below [49]. However, such a neuron would require a very high spontaneous firing rate to encode the full range of negative responses, which contrasts with experimental evidence that suggests low basal firing rates of layer 2/3 principal neurons [50,51]. Moreover, in a system where single neurons encode bi-directional errors, a postsynaptic neuron that receives error signals must have a mechanism to subtract out the basal firing rate of a presynaptic neuron. Given the discrete and non-linear dynamics of spiking neurons, which renders accurate approximation of synaptic transmission non-trivial, our attempts to implement bi-directional error coding with spiking neurons led to inaccurate propagation of prediction errors. Therefore, we separated the error unit into two subtypes, one coding positive and the other coding negative error (Fig 2B). The two units are complementary in propagating prediction errors to representation units in the next higher area. A positive error unit (Eq. 12) integrates bottom-up excitatory inputs from representation units within the same area (where *W_l,l_* = 1), 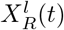, and top-down inhibitory inputs from representation units of the area above, 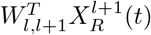, whereas a negative error unit (Eq. 13)has the opposite arrangement of excitatory-inhibitory synapses:

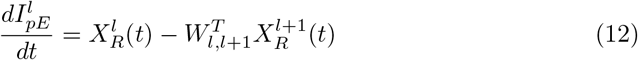

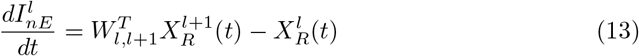

Note that bottom-up inputs to both positive and negative units are the same (i.e., *I_R^l^,pE^l^_* = *I_R^l^,nE^l^_*). Representation units and the two types of error unit within an area contain the same number of cells and connect to each other in a one-to-one fashion (i.e., *W_i,j_* = 1 where *i* = *j* and 0 elsewhere), so that error units receive the same bottom-up input, or its latent representation in the case of higher areas, and compare it to the top-down prediction.

#### Representation unit

Representation units infer the causes of sensory input via local interactions with error units in the area immediately below as well as those in the same area (Fig 2A). The interactions between two immediately adjacent areas are considered local, because they do not involve areas further down or up in the hierarchy (at least not directly) as would be commonly used in standard deep learning algorithms such as BP, which require global interactions from the top area to the lowest area. The inference process can be regarded as an iterative process of updating internal representations of sensory stimuli (or of neural activity of representation units, *I_R_^l^*), and is mathematically formalized as performing a gradient descent on the cost function of prediction error minimization with respect to the internal representation [16]. In SNN-PC, this first comes down to a sum of incoming synaptic current to each representation neuron:

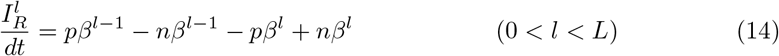

The first two terms in Eq. 14 represent the bottom-up positive and negative prediction error (*pβ*^*l*–1^ and *nβ*^*l*–1^), computed as weighted sums of NMDA currents arising from the connections between the two error units and representation units, respectively:

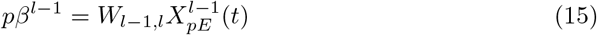

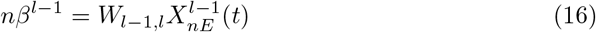

The last two elements of Eq. 14 are top-down positive and negative errors (*pβ^l^* and *nβ^l^*), respectively, which are connected to representation units in a one-to-one fashion:

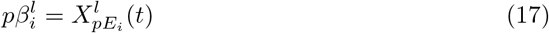

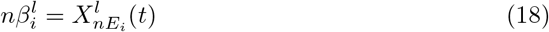

In the case of the highest area (area L or area 3 in Fig 3), these two terms are absent as it lacks top-down connections.

**Fig 3.**
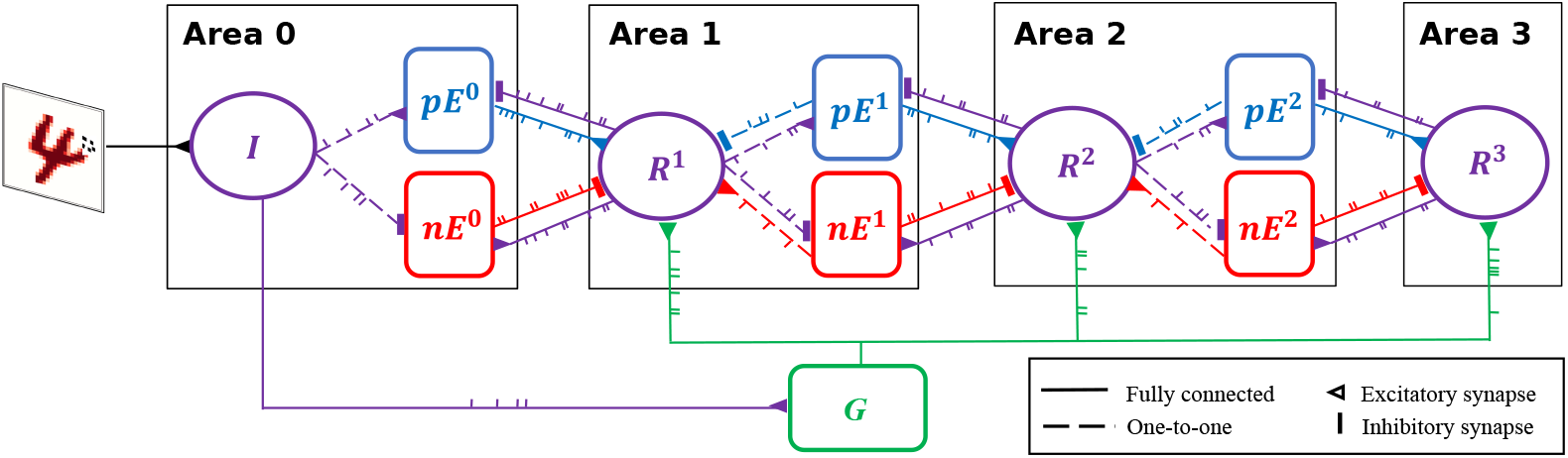
Spiking neural network for predictive coding (SNN-PC). SNN-PC modeling the first stages of the visual cortical processing hierarchy. The three areas roughly correspond to V1, V2, and V4, respectively (or LGN, V1, and V2). Each non-input area (*l* > 0) consists of a representation unit (purple circle; 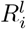) and two units (blue and red squares; 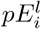 and 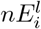) that encode positive and negative prediction error, respectively. The representation unit in Area 0 acts as an input unit (*R*^0^). Each pixel (the dotted box in the image of digit 4) is encoded by a single spiking neuron. Note that each unit consists of multiple spiking neurons. The feedforward gist pathway (*R*^0^ → *G* → *R^l^*) approximates the feedforward sweep of neuronal activity across the visual processing hierarchy. The solid lines between units indicate that they are fully connected, whereas the dotted lines indicate one-to-one connections. A triangle represents an excitatory synapse, whereas a thick vertical short ending represents an inhibitory synapse.

Sensory inputs are fed into the network via representation units in the lowest area (Area 0), each of which receives a constant current linearly proportional to the intensity of a pixel of the input image (Fig 3). The underlying assumption is that such a transduction is roughly comparable to a retinal image (with a resolution of a pixel). Given an MNIST digit sample (28 x 28 pixel image) as visual input, the number of units in Area 0 is 784.

### Feedforward gist pathway

Visual cortical processing can be parsed into two distinct processes [52]: 1) the fast feedforward sweep of neuronal activity across the visual processing hierarchy roughly within 150 ms of stimulus onset in primates, which is thought to generate coarse high-level representation of a visual scene and facilitates gist perception and rapid object recognition [40, 41, 53–55]; and 2) slow recurrent processing that iteratively refines the representation (a process henceforth referred to as inference). SNN-PC implements the former process with a FFG pathway and the latter process with the PC hierarchy.

The FFG pathway approximates the feedforward sweep across the visual hierarchy via sparse random projections from input to gist units (Fig 4):

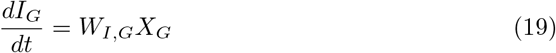

The weights between input and gist units (*W_I,G_* in Eq. 19) were randomly sampled from a Gaussian distribution, the mean and standard deviation of which were defined as a ratio between the number of pre- and postsynaptic units. To induce sparsity, the connection probability between input and gist units was set to a low value (*P_c_* = 0.05; Fig 4A).

**Fig 4.**
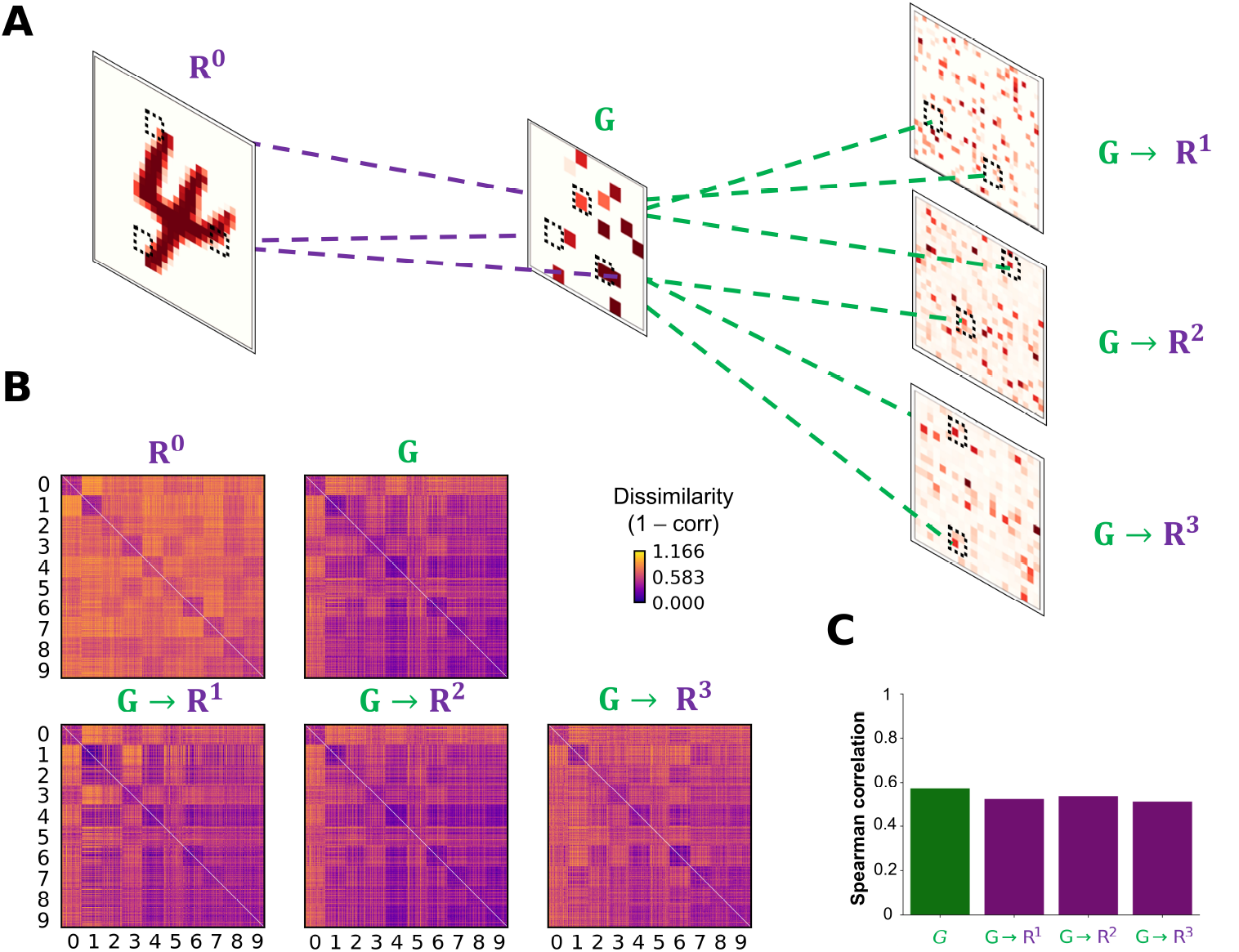
The fast feedforward gist pathway. (A) A fast feedforward sweep of neural activity from input (*R*^0^) to representation units (*R^l^* where *l* ∈ {1, 2, 3}) via gist units (*G*) is instantiated by sparse random connections to generate a spatially reduced, abstract representation of an input image, which provides an informative baseline for the upcoming iterations of recurrent processing. Purple dotted lines show example projections from input to gist units, whereas green dotted lines show example projections from gist to representation units in areas 1-3 of the visual processing hierarchy. Each tilted square represents an example neural activity pattern activated by an input image in each area. The colored elements in each area correspond to mean incoming synaptic currents. (B) A comparison between the representational dissimilarity matrix (RDM) of different areas involved in the feedforward gist pathway within the first 150 ms of simulation: the first row shows RDMs of mean synaptic currents into input (*R*^0^) and gist (*G*) units, whereas the second row shows mean synaptic currents from gist units to representation units in different areas (*G* → *R^l^*; *W_G,R^l^_ X_G_* where *l* ∈ {1, 2, 3}), excluding other synaptic currents from the recurrent dynamics of the PC hierarchy. Each element in an RDM represents a measure of dissimilarity between 100 different samples of digits (0-9). (C) Representational similarity between the RDM of input images (*R*^0^) to the rest of areas. Strong correlations indicate that the representational geometry of input images is retained across the FFG pathway (*R*^0^ → *G*) and that representation units in each cortical area (*G* → *R^l^* where *l* ∈ {1, 2, 3}) receive informative stimulus-based priors about the upcoming inference.

To reflect the increasing receptive field size and complexity of tuning properties when ascending the visual processing hierarchy, the number of gist units (144) is set to be smaller than input units (784). The resulting neuronal activity patterns in gist units therefore correspond to the coarse-grained representation of incoming sensory input. As all input images are processed by the same set of non-plastic, sparse random weights, images that share more features (e.g., two samples belonging to the category of digit ‘1’) have a higher chance to generate similar neural activity patterns in gist units than those that share less (e.g., a sample belonging to digit ‘0’ and another belonging to ‘1’); in other words, by statistically sampling the same area in the visual field given different images, the latent representations of input images in gist units retain the representational geometry of input images (Fig 4B-C).

Gist units then project to representation units in each area of the PC hierarchy to modulate their activity. The synaptic input coming from gist units can be implemented into the inference step (Eq. 14) by adding an extra term (*W_G,R^l^_ X^G^*):

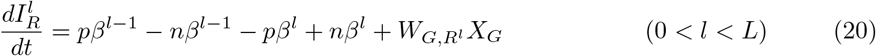

The FFG pathway runs in parallel with the PC hierarchy and lasts as long as the stimulus to provide a high-level, coarse representation of the incoming sensory input to representation units in each PC area (*W_G,R^l^_ X_G_*; Eq. 20) as a baseline activity.

In summary, the FFG pathway serves the role of initializing the neuronal activity of representation units in each area with a coarse representation of the incoming sensory input (e.g., the gist of a scene or object). Instead of starting the iterative process of prediction error minimization from zero or arbitrary activity in representation units, the gist-like latent representation of incoming sensory inputs operates in a biologically plausible manner.

### Rate-based Hebbian learning

With non-differentiable binary spike signals, *s*(*t*), the error gradient required to correct the internal model cannot be obtained. However, we can use exponentially filtered spike trains, *X*(*t*), to obtain the ingredients required for Hebbian learning (Fig 5).

**Fig 5.**
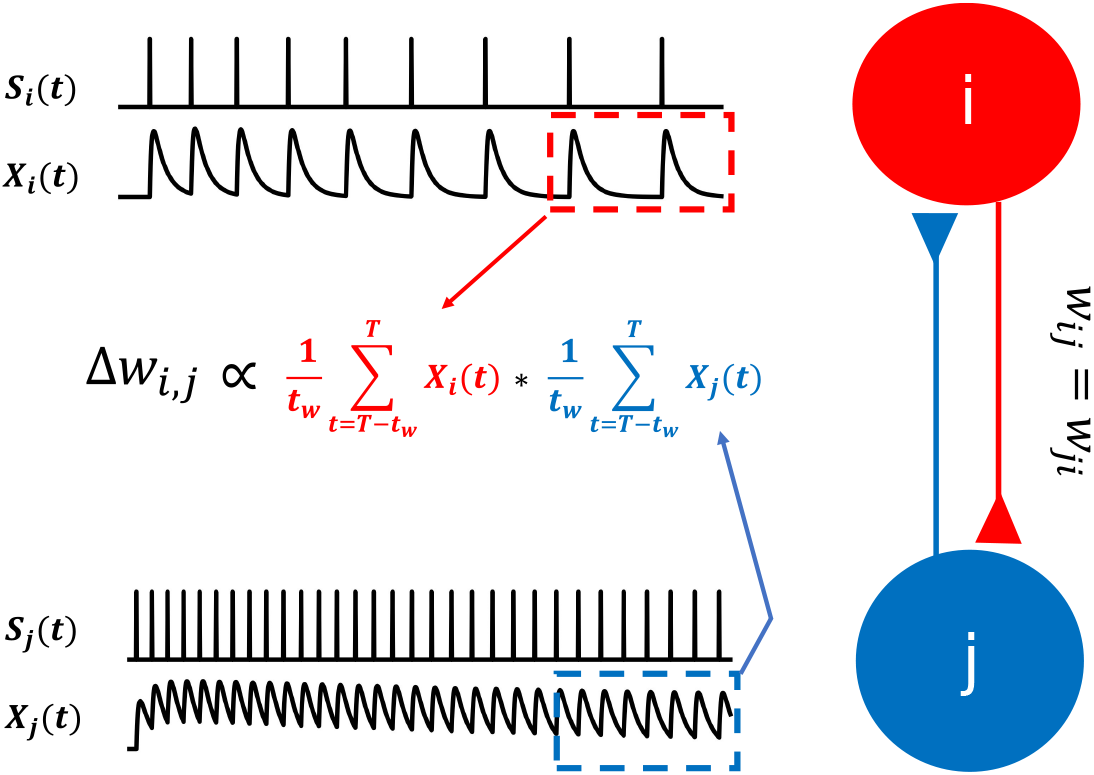
Hebbian learning for spiking neurons. In SNN-PC, synaptic plasticity is mediated by Hebbian learning. Instead of firing rates used in artificial neural networks, SNN-PC approximates NMDA receptor mediated postsynaptic calcium dynamics, *X_i_*(*t*) and *X_j_*(*t*), in each cell, based on incoming spike trains, *s_i_*(*t*) and *s_j_*(*t*), and leverages them to compute a biologically plausible learning gradient, Δ*w_ij_*. The red and blue dotted box indicate the time window, the last *t_w_* ms (from *T* – *t_w_* to *T*, where *T* is the total duration of stimulus presentation), during which the approximate postsynaptic calcium transient signals are averaged.

A weight matrix, *W*_*l,l*+1_, between error and representation units (Fig 3), is updated:

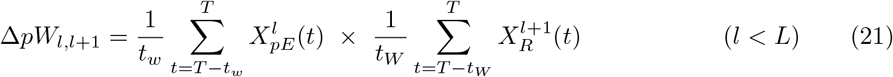

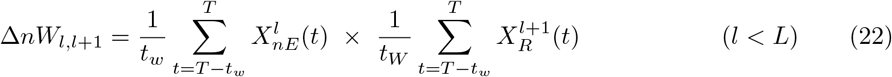

The two terms in Eq. 21 and 22 are mean NMDA current amplitudes entering the postsynaptic site of error and representation units from the last *t_W_* milliseconds (ms) of stimulus presentation (total duration = T ms): Eq. 21 specifies the use of positive error and Eq. 22 of negative error. Apart from convergence and stability purposes to accommodate spiking dynamics, taking this mean reflects the calcium dynamics in dendritic spines, which induce NMDA receptor-dependent long term plasticity and depression (LTP and LTD) [56–58]. The positive error units contact representation units with excitatory synapses to increase the calcium influx into representation units and can induce LTP, whereas the negative error units make inhibitory synaptic contact to decrease calcium influx and induce LTD [59]. Combining the two equations (Eq. 21 and 22) results in a weight update that is a linear combination of the Hebbian error gradients obtained between postsynaptic representation units and the two types of presynaptic error units:

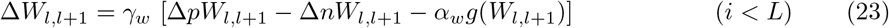

The weight change is controlled by the learning rate, *γ_w_*. The last term, *α_w_ g*(*W*_*l,l*+1_), models the passive decay of weights and imposes a Laplacian prior on the weights (i.e., L1 regularization) [35]:

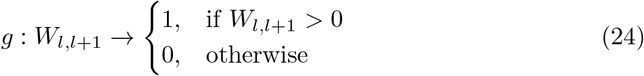

The resulting unsupervised learning algorithm can be regarded as Hebbian learning with biologically plausible learning signals, which requires only locally available information. Note that only inter-areal weights (*W*_*l,l*+1_) are subject to synaptic plasticity. The intra-areal weights (W?;) are fixed.

### Simulation details

#### Preprocessing of input image

The input images were normalized to unit vectors to limit the variance among pixel intensity distributions across different digits and samples and scaled to a range between 600 and 3000 *pA,* within which the input and output synaptic currents approximated a linear relationship.

##### Network size

Each area consisted of the same number of positive error units, negative error units, and representation units (Area 0 = 784 * 3 = 2352; Area 1 = 1296 * 3 = 3888; Area 2 = 1024 * 3 = 3072), except the top area (Area 3) that only contained 784 representation units (Table 2). There were 144 gist units. In total, the number of units in SNN-PC was 10240. Out of 6298176 total synapses in the network, 6291968 inter-areal synapses were subject to synaptic plasticity.

**Table 2.** Parameters for simulation

| Parameter | Meaning | Value |
| --- | --- | --- |
| $n_R^0$ | number of units in $R^0$ | 784 |
| $n_{pE}^0$ | number of units in $pE^0$ | 784 |
| $n_{nE}^0$ | number of units in $nE^0$ | 784 |
| $n_R^1$ | number of units in $R^1$ | 1296 |
| $n_{pE}^1$ | number of units in $pE^1$ | 1296 |
| $n_{nE}^1$ | number of units in $nE^1$ | 1296 |
| $n_R^2$ | number of units in $R^2$ | 1024 |
| $n_{pE}^2$ | number of units in $pE^2$ | 1024 |
| $n_{nE}^2$ | number of units in $nE^2$ | 1024 |
| $n_R^3$ | number of units in $R^3$ | 784 |
| $n_G$ | number of units in $G$ | 144 |
| $dt$ | simulation time step | 0.1 ms |
| $\tau_w$ | time window for synaptic plasticity | 100 ms |
| $T$ | total simulation time per sample | 350 ms |
| $\gamma_w$ | learning rate for synaptic plasticity | $1e^{-9}$ |
| $\alpha_w$ | regularizer for synaptic plasticity | $1e^{-5}$ |
| $n_{total}^{train}$ | total number of samples in training set | 10240 |
| $n_{total}^{test}$ | total number of samples in test set | 2560 |
| $n_{class}^{train}$ | number of samples per class in training set | 1024 |
| $n_{class}^{test}$ | number of samples per class in test set | 256 |
| $n_{batch}^{train}$ | number of mini-batches | 80 |
| $n_{sample}^{batch}$ | number of samples in a mini-batch | 128 |
| $n_{epoch}^{train}$ | number of training epochs | 60 |

##### Training and testing

In order to test whether SNN-PC can learn statistical regularities of incoming sensory inputs and build latent representations thereof, we trained SNN-PC with a subset of the MNIST handwritten digit image dataset. The training set (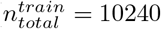; Table 2) consisted of many different image samples per image class (digits 0-9; 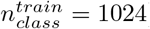). For efficient learning, we used mini-batch training 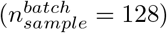. During a single training epoch, the network goes through all mini-batches 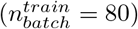. After each mini-batch, the synaptic weights are updated. After 60 training epochs 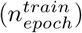, we tested the model to infer image samples it had not been exposed to during the training, taken from a test set 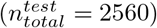.

##### Simulation

The solutions to differential equations for inference were numerically computed using Euler’s method with a temporal resolution of 0.1 ms (i.e., *dt* = 0.1 ms). Note that the internal dynamics of all units in SNN-PC (*V, a, X*, and *Y*) were updated concurrently at each time step. Each sample was presented to the network for 350 ms (i.e., *T* = 350ms) and repeated 60 times. Whenever a new image sample was presented, the internal dynamics of all units were reset to initial values (*V* = –70*mV*, and a = *X* = *Y* = 0).

For learning, we took the mean over the last 100 ms (250-350 ms) of synaptic current relative to the onset of the stimulus to ensure convergence. Weights were initialized randomly, but strictly positive, by sampling from a half-normal distribution around zero mean and a small standard deviation (0.3). For each weight update, a learning rate of 1e^-9^ and a regularization parameter of 1*e*^-5^ were used.

## Results

To test the perceptual capacity of SNN-PC, we trained it with a subset of the MNIST handwritten image dataset (n=10240; a full set has 60000 images) and tested whether it could learn to generate internal representations of novel images (n=2560) across the hierarchy, which can be used to reconstruct the original images.

The success of learning can first be qualitatively assessed from the error units in the lowest area (Area 0), which receive bottom-up sensory inputs that directly correspond to pixel intensities of input images and top-down predictions that reconstruct them. By organizing neural activity patterns that correspond to prediction (*W*_*R*_1_,*E*_0__ 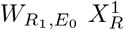; *Area* 1 → 0) into the input shape (28×28 pixels), we can visualize the generative capacity of SNN-PC, which can reconstruct novel images it had not seen during training (Fig 6A).

**Fig 6.**
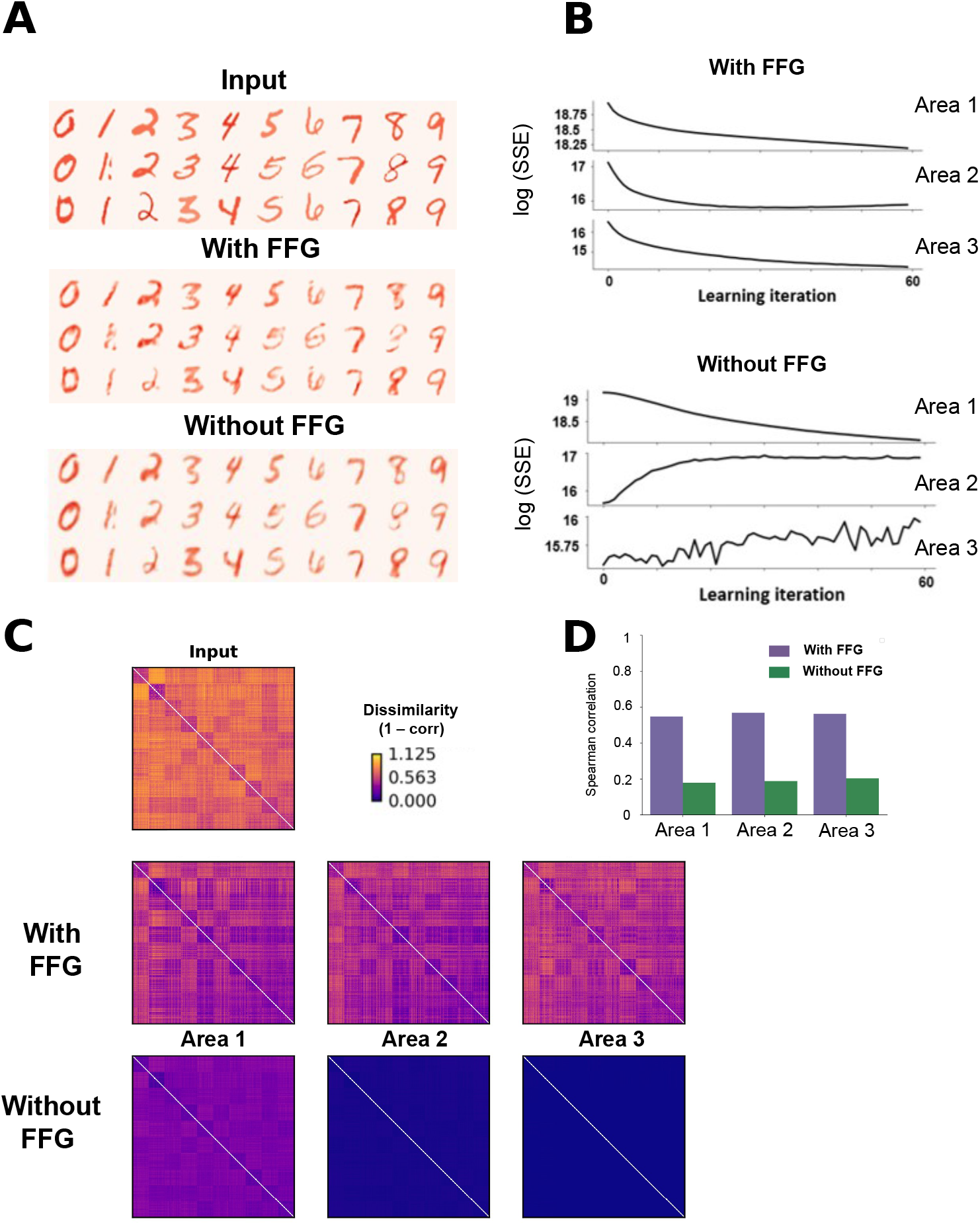
Learning of MNIST handwritten digits. (A) Reconstruction of novel sensory inputs. Input image (top) is compared to the prediction made from the area immediately above (Area 1) by SNN-PC with (middle) and without the FFG pathway (bottom). (B) Inference. Sum of squared prediction errors (SSE) in each area across training epochs computed by SNN-PC with (top) and without (bottom) the FFG pathway. The large values of prediction errors are due to their unit of measurement in pA and the summation over a large number of neurons (784 in Area 1, 1296 in Area 2, and 1024 in Area 3). An average prediction error per neuron in each area corresponds to 328, 83, and 34 pA, respectively. (C) Representational similarity analysis. Representational dissimilarity matrices constructed with test images in the input area (top) and inferred causes of those test images in each area from SNN-PC with (middle) and without (bottom) the FFG pathway. (D) Second-order representational similarity analysis. The RDM of test images in the input area is compared against the RDMs of inferred causes of those test images in each area from SNN-PC with (dark purple) and without (light purple) the FFG pathway.

Neural activity patterns that correspond to predictions in higher areas bear no immediate semantic meaning to a human beholder; therefore, they are called latent representations. To show that learning takes place in higher layers as well, we show the sum of squared errors (SSE), a quantitative measure of the discrepancy between incoming inputs and their predictions, in each area (Fig 6B). Note that the SSEs are displayed on a logarithmic scale, because prediction errors were measured in picoampere (pA) and summed over a large number of neurons. For the model with FFG pathway, the decreasing SSEs in all three areas across learning epochs indicate that SNN-PC can hierarchically extract statistical regularities from sensory inputs and update its internal model to more accurately infer their causes (top panel; Fig 6B).

To examine the internal representations the model developed of input images, we performed a representational similarity analysis (RSA) [60]. In each PC area (*R*^0^ - *R*^3^), all pairwise dissimilarities (1 - correlation) among the firing rate patterns corresponding to the test set of images were computed and organized into a matrix format (RDM; Fig 6C). The consistent pattern in RDMs of input images (*R*^0^) and internal representations across the PC hierarchy (*R*^1^ - *R*^3^; middle row; Fig 6C), as well as their strong correlations (purple bars; Fig 6D), show that the representational geometry of input images is retained in the processing hierarchy.

Note that the RDMs in Fig 6C compare firing rate patterns between input images, while the RDMs in Fig 4 compare mean synaptic current. In the former case, we demonstrate that the outgoing signals from representation units in each area (i.e., firing rates) preserve the representational geometry of input images, forming latent representations of the inputs and allowing for a consistent flow of information about the stimulus in the system. In the latter case, we show that FFG signals impinging on representation units (i.e., mean synaptic current from gist to representation units), independent of other signals caused by the recurrent dynamics of PC, retain the representational geometry of input images, providing an informative ’stimulus-based prior’ for the upcoming inference process.

We also trained and tested a model on the same task without the FFG pathway to analyze its effect on the perceptual capacity of SNN-PC. Instead of FFG inputs, representation units received random background noise. The SSE in the lowest area still decreased monotonically (Area 1 in the bottom panel; Fig 6B) without FFG inputs, leading to successful reconstructions of input images (bottom row; Fig 6A). However, the SSEs in higher areas increased (bottom panel; Fig 6B), which resulted in indistinguishable representations of sensory inputs in higher areas (bottom row; Fig 6C) that did not retain the representational geometry of input images. The representations of input images formed by higher areas were weakly correlated to input images (green bars; Fig 6D).

Furthermore, we examined the discriminative capacity of SNN-PC by training a linear regression classifier on neural activity patterns of representation units in response to a training set (n = 10240) and testing on a randomly sampled test set (n = 2560). The decoding accuracy was determined by the number of correct class predictions. For statistical comparison, namely Mann-Whitney’s U-test, the classifier was tested with 1000 different sets. Note that separate classifiers were used for a model with the FFG pathway and another without, to evaluate the effect of FFG inputs on the discriminative capacity of SNN-PC. As shown in Fig 7, neural activity patterns with FFG inputs showed significantly higher decoding accuracy than those without in all three areas (*p* < 0.001). Also, decoding accuracy decreased across the hierarchy when the model lacked the FFG pathway (*p* < 0.001). On the other hand, with the FFG pathway, the decoding accuracy was not significantly different in Area 1 and 2 but decreased significantly in Area 3 (p < 0.001).

**Fig 7.**
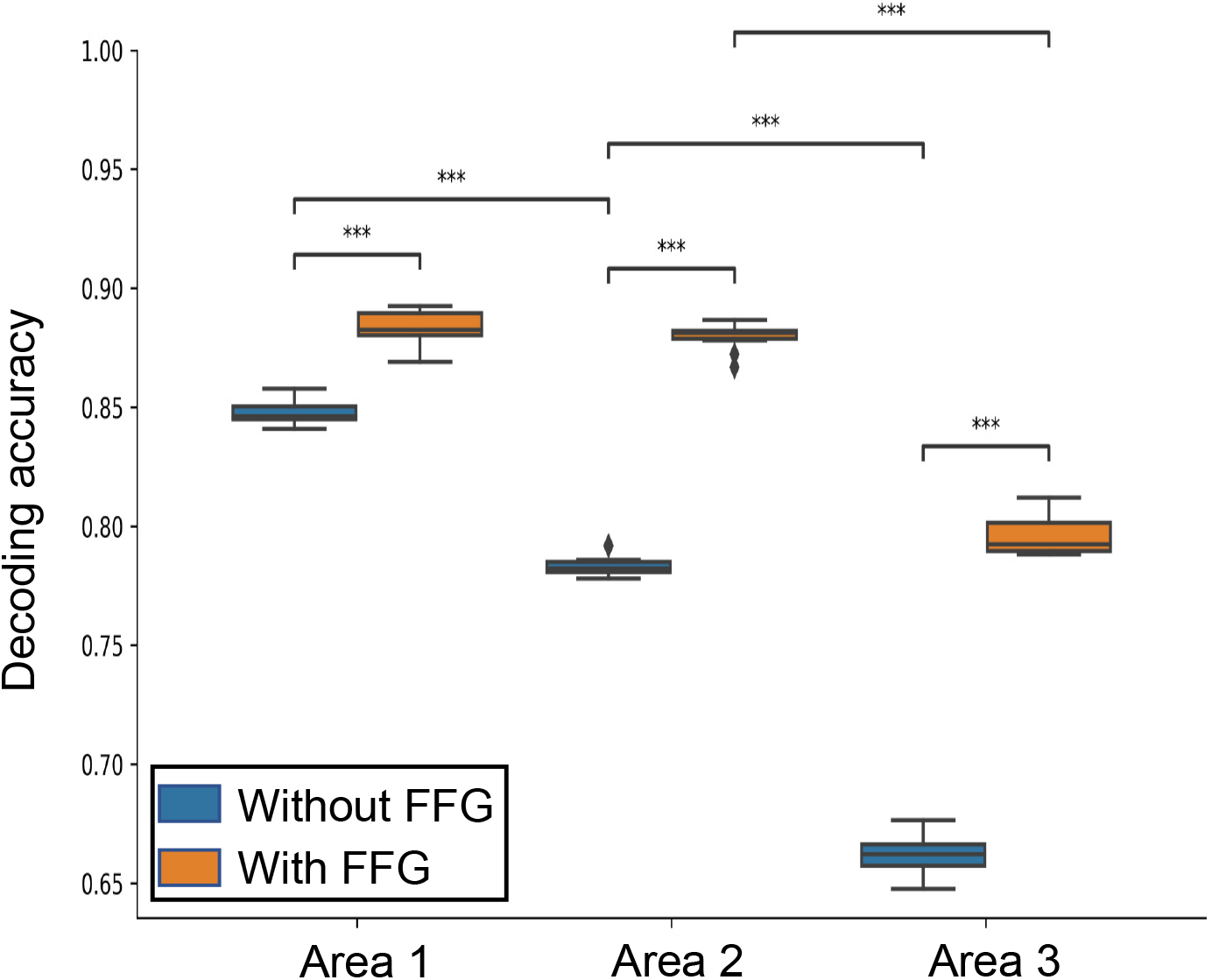
Digit classification performance. A boxplot showing accuracy of digit prediction decoded from internal representations of handwritten digit images in each PC area. Blue boxes represent performance of a classifier trained with patterns generated by a SNN-PC without the FFG pathway, whereas orange boxes show results of a classifier trained with patterns from a SNN-PC with the FFG pathway. Each box represents the interquartile range (*Q*_1_ - *Q*_3_) of the distribution of decoding accuracy on 1000 randomly sampled sets. Horizontal lines of the boxes denote the first, second, and third quartiles. Whiskers represent the minimum and the maximum of each accuracy distribution. Diamonds denote outliers. The statistical significance of difference between each group is noted with asterisks (***: *p* < 0.001).

## Discussion

We have described how to build a biologically grounded neural network for visual perception (SNN-PC), based on the following four components: 1) a predictive coding model that provides computational algorithms and a neural architecture for generative perceptual inference via recurrent sensory processing; 2) a FFG pathway that accounts for rapid feedforward processing in the visual cortical system; 3) spiking neurons that reflect the time-varying pulsatile behavior of neurons better than rate-based neurons; and 4) Hebbian learning that mediates synaptic plasticity. Our results show that a neural network model that combines PC and the fast feedforward sweep can perform perceptual inference and learning of two-dimensional visual images via neurobiologically grounded synaptic transmission and plasticity mechanisms. The model could also learn more complex images and scale up to include more areas (data not shown).

While inspired by biological neurons, artificial neural networks make many assumptions for the sake of functional and computational efficiency. For instance, the underlying assumption behind using firing rates as a measure of neural activity is that information is rate coded. However, the brain is also thought to encode information using both the timing of spikes (e.g., phase coding and neural synchrony [61–64]) and aggregate responses of neuronal ensembles (i.e., population coding [65–68]). By collapsing the temporal dimension into an arbitrary time window, artificial neurons cannot leverage asynchronous, event-based, and sparse information processing for energy efficiency [69–71]. Moreover, the non-local, end-to-end error propagation of BP poses a serious challenge to biological plausibility [72–76]. To create a system that performs complex cognitive behaviors on par with a human agent, we need to study and incorporate principles of neural computation and architectures from the biological agent we want to mimic, namely the mammalian brain. A straightforward choice in pursuing such endeavors therefore was to introduce spiking neurons, as they provide a biophysically realistic level of detail to simulate basic computations in the brain.

The model learned to reconstruct and develop latent representations of the MNIST handwritten digit images using only a small subset of the image dataset (1/6 of the full dataset) and an unsupervised learning method that requires only locally available information at each level of the hierarchy. The remaining errors below the convergence points in each panel of Fig 6B probably had little effect on inference and learning performance, because the average prediction error per neuron (Area 1 ≈ 328 pA, Area 2 ≈ 83 pA, and Area 3 ≈ 34 pA) falls below the firing threshold of spiking neurons used in SNN-PC in all areas. Nonetheless, these residual errors must reflect the noisy approximation of learning signals. In SNN-PC, the exponential low-pass filtering of discrete spike trains from a source unit generates a continuous signal that approximates the effect of presynaptic spikes targeting the postsynaptic membrane potential (i.e., calcium transient in spines) online. On the other hand, the strength of synaptic change is determined at discrete time points by taking the mean of the calcium transient over a short time interval (100 ms). Performing gradient descent with a point approximation (i.e., taking the mean of incoming synaptic currents) of an asynchronous and non-linear signal to drive its trajectory to a stationary point cannot yield a precise solution.

Furthermore, based on what it had learned, SNN-PC was able to generalize to novel images. Despite PC being a generative model, the objective function of which does not optimize a discriminative problem, we were able to decode class labels from neural activity patterns with a high accuracy in early areas (≈ 90% in Area 1 and 2; Fig 7). Without any explicit network design such as one implementing receptive fields or convolutional layers, the invariant latent representation of digit classes naturally emerged during both training and test phases. However, even with a FFG pathway, the highest area (Area 3) showed a significant drop in decoding accuracy (*p* < 0.0001). This is most likely due to the lack of top-down inputs in Area 3, in contrast to the middle areas (Area 1 and 2) where they are present and provide guidance for inference.

### Novel features of the predictive coding model with spiking neurons

To our knowledge, no predictive coding model has been proposed before that is operating purely with spiking neurons, except for the spiking neural coding network (SpNCN) proposed in [77]. While SNN-PC and SpNCN similarly implement synaptic transmission (i.e., low-pass filtering of spike trains) and weight updating (i.e., Hebbian learning), SNN-PC takes into account the following additional constraints. First, spiking neurons in SNN-PC are based on the AdEx model, which offers a biophysically more accurate description of a neuron’s behavior than the leaky-and-integrate fire (LIF) model used in SpNCN [42].

A second novel feature of SNN-PC is that it has two types of error unit to compute positive and negative error separately to avoid employing unrealistic negative synaptic weights for inhibitory connections (Fig 2B). Despite having solved the implausible negative weight problem, all units in SNN-PC (when viewed as single neurons) do not adhere to another biological reality (i.e., Dale’s principle) as they form both excitatory (in case of the *R^l^* → *pE*^*l*–1^ projection) and inhibitory (*R^l^* → *nE*^*l*–1^) synapses onto other units. However, such violation can be mitigated by replacing computational units in SNN-PC (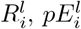, and 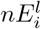; Fig 2B) by cortical microcircuits that consist of pyramidal cells and interneurons (e.g., green triangle and orange circle wrapped inside purple, blue, and red contours, respectively; Fig 2C). Note that we used one pyramidal cell and one interneuron in a microcircuit for visual presentation purpose only. Using this microcircuit scheme, we predict that an excitatory synapse between two microcircuits (e.g., 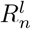 and 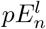) is formed between their pyramidal cells, whereas an inhibitory synapse between the two microcircuits consists of an excitatory connection from pyramidal cells in a microcircuit (e.g., 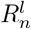) to interneurons in another microcircuit (e.g., 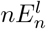), which then inhibits pyramidal cells in the same microcircuit (e.g., 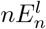). Future research will have to show how PC can be implemented using known anatomical connections [78], laminar organization [10], and different cell types (e.g., pyramidal, SST, VIP, and PV; [79]).

Besides offering a biologically plausible solution to accommodate binary spiking dynamics, the explicit separation of error units into positive and negative elements may also be beneficial for the algorithmic and computational efficiency of neuromorphic hardware. It only requires a straightforward subtraction between input and prediction, whereas bi-directional error units (such as postulated in reinforcement learning models based on prediction error coding by mesencephalic dopamine neurons [49]) must first compute the difference and then compare it against a basal firing rate to determine the sign of the error. With a certain basal firing rate, positive and negative errors can be encoded by the range above and below it, respectively. However, for a full coverage of prediction error ranges, bidirectional error units must maintain a high basal firing rate, thereby leading to a higher energy cost. The neuron targeted by a bidirectional error unit would also have to be equipped with a mechanism to discount for the basal firing rate to obtain the true prediction error.

### Feedforward gist

A third novel feature of SNN-PC is the FFG pathway, which combines the fast feedforward sweep and the slow recurrent PC to account for a more comprehensive picture of visual processing. PC reconciles bottom-up and top-down accounts of perception by casting it as an inferential process that involves hierarchical recurrent interactions. However, the inference process requires multiple loops of recurrent processing to converge on an accurate representation of incoming sensory input. The expected latency of visual responses arising from recurrent predictive processing is not in accordance with the rapid forward spread of object- and context-sensitive neuronal activity across the visual cortical hierarchy within 100 ms of stimulus onset [52]. While the precise contributions of fast feedforward and slow recurrent processes to perception are yet to be determined [80], we aimed to combine these two temporally distinct processes by integrating the FFG pathway in a PC architecture and whether the FFG pathway can improve network performance.

Reflecting the temporal dichotomy of the two visual processes, the FFG pathway quickly establishes a high-level, coarse representation of input signals (e.g., the gist of a scene or object) and feeds it to each area of the PC hierarchy to aid the recurrent processing that slowly refines the representation. Instead of starting the iterative process of prediction error minimization from zero or arbitrary activity in representation units, the gist-like latent representation of incoming sensory inputs offers a biologically plausible starting point for the recurrent processing. This suggests a novel function of the feedforward activity relative to the classic hypothesis of rapid image recognition [39].

When we tested the impact of the FFG pathway on the learning of two-dimensional visual images, our results showed that prediction errors decrease across training epochs in all areas when the FFG pathway was present (with FFG; Fig 6B), whereas in its absence, SSE converged only in Area 1 (without FFG;Fig 6B). The main reason for this disparity can be attributed to the difference in the nature of synaptic inputs given to the representation units in each area prior to the moment at which the recurrent dynamics of PC initialized their neuronal activity. In the model with a FFG pathway, these pre-PC inputs are consistent across the PC hierarchy, as they come from a single source (gist units) that is related to the image. This helps the development of consistent representations across the hierarchy, leading to convergence in all areas. On the other hand, in the model without a FFG pathway, the pre-PC inputs were independently sampled Gaussian noise, making them inconsistent across the hierarchy. Given the local processing property of PC, each area initiated its inference at random places along the slope of the error gradient, independent of each other; on contrary, these spots will be closer to each other in the model with the FFG pathway. Such inconsistency in the pre-PC inputs likely hindered the development of consistent representations across the hierarchy. Nonetheless, the bottom-most area (Area 1) still converged, owing to the constant input directly corresponding to the stimulus, leading to good reconstruction of input images (without FFG; Fig 6A). On the contrary, higher areas (Area 2 and 3) received bottom-up input that fluctuates over time due to ongoing inference process at each level of the hierarchy. These findings suggest that the FFG plays a crucial role in the development of consistent representations across the cortical hierarchy by placing an a priori constraint on the inference process.

Meanwhile, it is not clear whether and how the same populations of neurons are involved in both the feedforward sweep and recurrent PC. Instead of performing a series of feedforward feature extraction and integration steps to elicit object-sensitive responses in high visual areas, feedforward connections in the PC network convey prediction errors. Therefore, the two processes might take separate routes. For instance, recurrent processes governed by PC may occur via cortico-cortical pathways, whereas the feedforward sweep may be mediated by the pulvino-cortical pathway [81]. Alternatively, the FFG pathway can be explained as a combination of the fast feedforward sweep and subsequent top-down modulation: an instantaneous feedforward sweep activates gist units, conceptualized as IT, PFC, or other high-level cells; this is followed by top-down projections from gist to representation units in lower visual areas (e.g., V1 to V4).

In both scenarios, we assume innate and non-plastic feedforward connections. A recent study [82] examined how such feedforward connections can be trained via amortized inference and also showed robust perceptual capacity with shorter error convergence time and fewer training samples. However, we note that SNN-PC is spike-based and unsupervised, whereas the model in [82] is rate-based and uses a mix of supervised and unsupervised learning.

### Future directions

There are various ways in which future studies may extend the biological details and/or perceptual capacities of SNN-PC. To name a few, first, a point spiking neuron can be replaced by a cortical microcircuit with multiple interneurons (e.g., Fig 2C) with a columnar organization to replicate experimental findings and make predictions for new experiments. Second, spiking neuron behavior or connectivity can be altered to implement receptive fields and response properties to construct a invariant object representation. Third, self-recurrent loops or online learning rules such as STDP may be employed to deal with a continuous stream of sensory inputs. Fourth, a population coding regime can be implemented to improve the reliability of signal transmission [83]. Fifth, a different sensory modality can be added to perform multi-sensory integration, following the rate-based predecessor of our current SNN-PC model [35], which has been implemented in a rodent robot that performs bimodal integration of vision and touch to navigate in a maze [84]. Sixth, the network can be scaled up to better reflect the areas involved in the visual processing hierarchy.

## Conclusion

We have described how to build a PC model of visual perception using biologically plausible components such as spiking neurons, Hebbian learning, and a FFG pathway. While SNN-PC is far from reaching the cognitive performance of state-of-the-art artificial intelligence or capturing the detailed biological mechanisms underlying perception, it is one of the first purely spike-based and completely unsupervised PC models of visual perception. As such it successfully performs perceptual inference and learning, and thereby may inspire machine learning, neuromorphics, neuroscience and cognitive science communities to seek avenues moving closer to mimic the nature’s most intelligent and efficient system, the brain.

## Acknowledgments

The authors would like to thank Matthias Brucklacher, Giulia Moreni, Giovanni Pezzulo, and Walter Senn for constructive discussions.

## References

1. Pizlo Z. Perception viewed as an inverse problem. Vision research. 2001;41(24):3145–3161.

2. Spratling MW. A review of predictive coding algorithms. Brain and cognition. 2017;112:92–97.

3. Fechner GT. Elements of psychophysics, 1860. Appleton-Century-Crofts; 1948.

4. Kant I. Critique of pure reason. 1781. Modern Classical Philosophers, Cambridge, MA: Houghton Mifflin. 1908; p. 370–456.

5. Von Helmholtz H. Treatise on physiological optics. vol. 3. Courier Corporation; 2013.

6. Gregory RL. The Intelligent Eye. Mcgraw-Hill; 1970.

7. MacKay DM. In: Shannon CE, McCarthy J, editors. The Epistemological Problem for Automata. Princeton University Press; 2016. p. 235–252. Available from: https://doi.org/10.1515/9781400882618-012.

8. Neisser U. Cognitive psychology / Ulric Neisser. Appleton-Century-Crofts New York; 1967.

9. Pennartz C. The brain’s representational power: on consciousness and the integration of modalities. Cambridge, Massachusetts London, England: The MIT Press; 2015.

10. Bastos AM, Usrey WM, Adams RA, Mangun GR, Fries P, Friston KJ. Canonical microcircuits for predictive coding. Neuron. 2012;76(4):695–711.

11. Felleman DJ, Van Essen DC. Distributed hierarchical processing in the primate cerebral cortex. Cerebral cortex (New York, NY: 1991). 1991;1(1):1–47.

12. Keller GB, Bonhoeffer T, Hübener M. Sensorimotor mismatch signals in primary visual cortex of the behaving mouse. Neuron. 2012;74(5):809–815.

13. Walsh KS, McGovern DP, Clark A, O’Connell RG. Evaluating the neurophysiological evidence for predictive processing as a model of perception. Annals of the new York Academy of Sciences. 2020;1464(1):242–268.

14. Srinivasan MV, Laughlin SB, Dubs A. Predictive coding: a fresh view of inhibition in the retina. Proceedings of the Royal Society of London Series B Biological Sciences. 1982;216(1205):427–459.

15. Mumford D. On the computational architecture of the neocortex. Biological cybernetics. 1992;66(3):241–251.

16. Rao RP, Ballard DH. Predictive coding in the visual cortex: a functional interpretation of some extra-classical receptive-field effects. Nature Neuroscience. 1999;2:79–87.

17. Friston K. A theory of cortical responses. Philosophical transactions of the Royal Society B: Biological sciences. 2005;360(1456):815–836.

18. Pennartz CM, Dora S, Muckli L, Lorteije JA. Towards a unified view on pathways and functions of neural recurrent processing. Trends in neurosciences. 2019;42(9):589–603.

19. Shannon CE. A mathematical theory of communication. The Bell System Technical Journal. 1948;27(3):379–423. doi: 10.1002/j.1538-7305.1948.tb01338.x.

20. Barlow H. Possible Principles Underlying the Transformations of Sensory Messages. Sensory Communication. 1961;1. doi: 10.7551/mitpress/9780262518420.003.0013.

21. Friston K. The free-energy principle: a unified brain theory? Nature reviews neuroscience. 2010;11(2):127–138.

22. Knill DC, Pouget A. The Bayesian brain: the role of uncertainty in neural coding and computation. Trends in Neurosciences. 2004;27(12):712–719.

23. Dayan P, Hinton GE, Neal RM, Zemel RS. The Helmholtz machine. Neural computation. 1995;7(5):889–904.

24. Hosoya T, Baccus SA, Meister M. Dynamic predictive coding by the retina. Nature. 2005;436(7047):71–77.

25. Jehee JF, Rothkopf C, Beck JM, Ballard DH. Learning receptive fields using predictive feedback. Journal of Physiology-Paris. 2006;100(1-3):125–132.

26. Spratling MW. Predictive coding as a model of response properties in cortical area V1. Journal of neuroscience. 2010;30(9):3531–3543.

27. Huang Y, Rao RP. Predictive coding. Wiley Interdisciplinary Reviews: Cognitive Science. 2011;2(5):580–593.

28. Wacongne C, Changeux JP, Dehaene S. A neuronal model of predictive coding accounting for the mismatch negativity. Journal of Neuroscience. 2012;32(11):3665–3678.

29. Spratling MW. A neural implementation of Bayesian inference based on predictive coding. Connection Science. 2016;28(4):346–383. doi: 10.1080/09540091.2016.1243655.

30. Lotter W, Kreiman G, Cox D. Deep Predictive Coding Networks for Video Prediction and Unsupervised Learning; 2016. Available from: https://arxiv.org/abs/1605.08104.

31. Whittington JCR, Bogacz R. An Approximation of the Error Backpropagation Algorithm in a Predictive Coding Network with Local Hebbian Synaptic Plasticity. Neural Computation. 2017;29(5):1229–1262. doi: 10.1162/NECO_a_00949.

32. Wen H, Han K, Shi J, Zhang Y, Culurciello E, Liu Z. Deep Predictive Coding Network for Object Recognition; 2018. Available from: https://arxiv.org/abs/1802.04762.

33. Van den Oord A, Li Y, Vinyals O. Representation learning with contrastive predictive coding. arXiv e-prints. 2018; p. arXiv–1807.

34. Han T, Xie W, Zisserman A. Video Representation Learning by Dense Predictive Coding; 2019. Available from: https://arxiv.org/abs/1909.04656.

35. Dora S, Bohte SM, Pennartz CMA. Deep Gated Hebbian Predictive Coding Accounts for Emergence of Complex Neural Response Properties Along the Visual Cortical Hierarchy. Frontiers in Computational Neuroscience. 2021;15:65.

36. Maass W. Networks of spiking neurons: the third generation of neural network models. Neural networks. 1997;10(9):1659–1671.

37. Gerstner W. Spiking neuron models: single neurons, populations, plasticity. Cambridge, U.K. New York: Cambridge University Press; 2002.

38. Oliva A, Torralba A. Building the gist of a scene: The role of global image features in recognition. Progress in brain research. 2006;155:23–36.

39. Thorpe S, Fize D, Marlot C. Speed of processing in the human visual system. nature. 1996;381(6582):520–522.

40. Serre T, Oliva A, Poggio T. A feedforward architecture accounts for rapid categorization. Proceedings of the National Academy of Sciences. 2007;104(15):6424–6429. doi: 10.1073/pnas.0700622104.

41. VanRullen R. The power of the feed-forward sweep. Advances in Cognitive Psychology. 2007;3(1-2):167.

42. Brette R, Gerstner W. Adaptive exponential integrate-and-fire model as an effective description of neuronal activity. Journal of neurophysiology. 2005;94(5):3637–3642.

43. Badel L, Lefort S, Brette R, Petersen CC, Gerstner W, Richardson MJ. Dynamic IV curves are reliable predictors of naturalistic pyramidal-neuron voltage traces. Journal of Neurophysiology. 2008;99(2):656–666.

44. Gerstner W, Kistler WM, Naud R, Paninski L. Neuronal dynamics. Cambridge, England: Cambridge University Press; 2014.

45. Forsythe ID, Westbrook GL. Slow excitatory postsynaptic currents mediated by N-methyl-D-aspartate receptors on cultured mouse central neurones. The Journal of Physiology. 1988;396(1):515–533.

46. McBain CJ, Dingledine R. Heterogeneity of synaptic glutamate receptors on CA3 stratum radiatum interneurones of rat hippocampus. The Journal of physiology. 1993;462(1):373–392.

47. Barria A, Malinow R. NMDA receptor subunit composition controls synaptic plasticity by regulating binding to CaMKII. Neuron. 2005;48(2):289–301.

48. Granger AJ, Nicoll RA. Expression mechanisms underlying long-term potentiation: a postsynaptic view, 10 years on. Philosophical Transactions of the Royal Society B: Biological Sciences. 2014;369(1633):20130136.

49. Schultz W, Dayan P, Montague PR. A Neural Substrate of Prediction and Reward. Science. 1997;275(5306):1593–1599. doi: 10.1126/science.275.5306.1593.

50. De Kock CPJ, Bruno RM, Spors H, Sakmann B. Layer- and cell-type-specific suprathreshold stimulus representation in rat primary somatosensory cortex. The Journal of Physiology. 2007;581(1):139–154. doi: https://doi.org/10.1113/jphysiol.2006.124321.

51. Perrenoud Q, Pennartz CM, Gentet LJ. Membrane potential dynamics of spontaneous and visually evoked gamma activity in V1 of awake mice. PLoS biology. 2016;14(2):21.

52. Lamme VA, Roelfsema PR. The distinct modes of vision offered by feedforward and recurrent processing. Trends in neurosciences. 2000;23(11):571–579.

53. Rousselet G, Joubert O, Fabre-Thorpe M. How long to get to the “gist” of real-world natural scenes? Visual cognition. 2005;12(6):852–877.

54. Liu H, Agam Y, Madsen JR, Kreiman G. Timing, timing, timing: fast decoding of object information from intracranial field potentials in human visual cortex. Neuron. 2009;62(2):281–290.

55. Cauchoix M, Crouzet SM, Fize D, Serre T. Fast ventral stream neural activity enables rapid visual categorization. NeuroImage. 2016;125:280–290.

56. Collingridge GL, Bliss T. NMDA receptors-their role in long-term potentiation. Trends in neurosciences. 1987;10(7):288–293.

57. Malenka RC, Nicoll RA. NMDA-receptor-dependent synaptic plasticity: multiple forms and mechanisms. Trends in neurosciences. 1993;16(12):521–527.

58. Lüscher C, Nicoll RA, Malenka RC, Muller D. Synaptic plasticity and dynamic modulation of the postsynaptic membrane. Nature neuroscience. 2000;3(6):545–550.

59. Mulkey RM, Malenka RC. Mechanisms underlying induction of homosynaptic long-term depression in area CA1 of the hippocampus. Neuron. 1992;9(5):967–975.

60. Kriegeskorte N, Mur M, Bandettini PA. Representational similarity analysis-connecting the branches of systems neuroscience. Frontiers in systems neuroscience. 2008;2:4.

61. Gray CM, König P, Engel AK, Singer W. Oscillatory responses in cat visual cortex exhibit inter-columnar synchronization which reflects global stimulus properties. Nature. 1989;338(6213):334–337.

62. O’Keefe J, Recce ML. Phase relationship between hippocampal place units and the EEG theta rhythm. Hippocampus. 1993;3(3):317–330.

63. Singer W. Neuronal synchrony: a versatile code for the definition of relations? Neuron. 1999;24(1):49–65.

64. Brette R. Computing with neural synchrony. PLoS computational biology. 2012;8(6):e1002561.

65. Georgopoulos AP, Schwartz AB, Kettner RE. Neuronal population coding of movement direction. Science. 1986;233(4771):1416–1419.

66. Lee C, Rohrer WH, Sparks DL. Population coding of saccadic eye movements by neurons in the superior colliculus. Nature. 1988;332(6162):357–360.

67. Averbeck BB, Latham PE, Pouget A. Neural correlations, population coding and computation. Nature reviews neuroscience. 2006;7(5):358–366.

68. Pouget A, Dayan P, Zemel R. Information processing with population codes. Nature Reviews Neuroscience. 2000;1(2):125–132.

69. Pfeiffer M, Pfeil T. Deep learning with spiking neurons: opportunities and challenges. Frontiers in neuroscience. 2018; p. 774.

70. Tavanaei A, Ghodrati M, Kheradpisheh SR, Masquelier T, Maida A. Deep learning in spiking neural networks. Neural networks. 2019;111:47–63.

71. Deng L, Wu Y, Hu X, Liang L, Ding Y, Li G, et al. Rethinking the performance comparison between SNNS and ANNS. Neural networks. 2020;121:294–307.

72. Rumelhart DE, Hinton GE, Williams RJ. Learning representations by back-propagating errors. nature. 1986;323(6088):533–536.

73. Bengio Y, Lee DH, Bornschein J, Mesnard T, Lin Z. Towards biologically plausible deep learning. arXiv preprint arXiv:150204156. 2015;.

74. Whittington JC, Bogacz R. Theories of error back-propagation in the brain. Trends in cognitive sciences. 2019;23(3):235–250.

75. Lillicrap TP, Santoro A, Marris L, Akerman CJ, Hinton G. Backpropagation and the brain. Nature Reviews Neuroscience. 2020;21(6):335–346.

76. Song Y, Lukasiewicz T, Xu Z, Bogacz R. Can the Brain Do Backpropagation?—Exact Implementation of Backpropagation in Predictive Coding Networks. Advances in neural information processing systems. 2020;33:22566–22579.

77. Ororbia A. Spiking Neural Predictive Coding for Continual Learning from Data Streams; 2019. Available from: https://arxiv.org/abs/1908.08655.

78. Douglas RJ, Martin K. A functional microcircuit for cat visual cortex. The Journal of physiology. 1991;440(1):735–769.

79. Keller GB, Mrsic-Flogel TD. Predictive Processing: A Canonical Cortical Computation. Neuron. 2018;100(2):424–435. doi: https://doi.org/10.1016/j.neuron.2018.10.003.

80. Kreiman G, Serre T. Beyond the feedforward sweep: feedback computations in the visual cortex. Annals of the New York Academy of Sciences. 2020;1464(1):222–241.

81. Jaramillo J, Mejias JF, Wang XJ. Engagement of pulvino-cortical feedforward and feedback pathways in cognitive computations. Neuron. 2019;101(2):321–336.

82. Tschantz A, Millidge B, Seth AK, Buckley CL. Hybrid Predictive Coding: Inferring, Fast and Slow. arXiv preprint arXiv:220402169. 2022;.

83. Boerlin M, Denève S. Spike-based population coding and working memory. PLoS computational biology. 2011;7(2):e1001080.

84. Pearson MJ, Dora S, Struckmeier O, Knowles TC, Mitchinson B, Tiwari K, et al. Multimodal representation learning for place recognition using deep Hebbian predictive coding. Frontiers in Robotics and AI. 2021;8.

